# Coordinated regulation of Citron kinase by CDK1 and Aurora B regulates midbody formation and stability

**DOI:** 10.1101/2025.08.25.672096

**Authors:** Luisa Capalbo, Ella F.J. Halcrow, Zuni I. Bassi, Pier Paolo D’Avino

**Affiliations:** Department of Pathology, University of Cambridge, Tennis Court Road, Cambridge, CB2 1QP, UK

**Keywords:** abscission, cell division, cytokinesis, midbody, phosphorylation

## Abstract

Many cell division events are regulated by protein phosphorylation, which can result from cross-talk mechanisms among mitotic kinases and phosphatases that have yet to be fully elucidated. Here, we report the characterization of a novel cross-talk mechanism by which CDK1 and Aurora B (AURKB) kinases regulate the distribution and interactions of Citron kinase (CIT-K). We show that CDK1 and AURKB phosphorylate two serine residues, S440 and S699, located adjacent to or within CIT-K coiled coil domain. S440 and S699 temporal phosphorylation profiles reflect the activity of the kinases responsible for their phosphorylation. Functional analyses using phospho mutants indicate that S699 phosphorylation is important for CIT-K localization and successful cytokinesis, while perturbing S440 phosphorylation leads to abnormal midbody formation and accumulation of post-mitotic midbody remnants (MBRs). Furthermore, we found that phosphorylation at either residue reduces the ability of CIT-K to interact with its midbody partners AURKB, KIF14 and KIF23/MKLP1. Together, our findings indicate that phosphorylation of CIT-K by CDK1 and AURKB regulates midbody formation and MBR stability by controlling the association of CIT-K with its partners. They expand our understanding of the mechanisms that regulate abscission and can lead to further insights into the role of MBRs in post-mitotic events.

## Introduction

Cell division is responsible for the correct partitioning of genomic and cytoplasmic contents between the two nascent daughter cells. The entire mitotic process is regulated by finely tuned signals that control the activity, localization, function, and interactions of proteins and protein complexes (Wieser and Pines, 2015). Due to the rapidity of mitotic events and chromatin inaccessibility, most mitotic processes are regulated by reversible post translational modifications, such as phosphorylation. The changes in the phosphorylation status of mitotic proteins are controlled by opposing enzymes, kinases and phosphatases, and these regulate every mitotic event, including entry into mitosis, the assembly of the mitotic spindle, chromosome alignment and segregation, and daughter cell separation during cytokinesis (Cuijpers and Vertegaal, 2018; Gelens et al., 2018; Holder et al., 2019; Nasa and Kettenbach, 2018). Although evidence of cross-talk mechanisms among different mitotic kinases and phosphatases exists (Cuijpers and Vertegaal, 2018), our understanding of the complexity and scope of these cross-talks is rather limited.

The last phase of cell division, cytokinesis, represents a powerful temporal window to study phosphorylation for various reasons. First, anaphase and cytokinesis are triggered by the degradation of cyclin B and consequent inactivation of its cyclin-dependent kinase 1 (CDK1) partner. This drop in CDK1 activity is complemented by increased activity of PP1 and PP2A serine/threonine phosphatases as well as by changes in the localization of other mitotic kinases, including Aurora B (AURKB), Citron kinase (CIT-K), and Polo-like kinase 1 (Plk1) (D’Avino and Capalbo, 2016; Holder et al., 2019). Together, these events lead to changes in the phosphorylation and activity of many proteins that drive a complete re-organization of the cytoskeleton (D’Avino et al., 2015). First, cells establish the position of the division plane through signals emanating from the spindle microtubules (MTs), which are re-organized into an array of antiparallel and interdigitating MTs, known as the central spindle, during anaphase. Spindle MTs also promote furrow ingression through the assembly and constriction of an actomyosin contractile ring, which compacts the central spindle and leave the daughter cells connected by an intercellular bridge in late cytokinesis. An electron and phase dense structure, first described by Flemming more than a century ago (Flemming, 1891), forms at the center of the intercellular bridge (Guizetti and Gerlich, 2010). This organelle, the midbody, is composed of an assortment of proteins with diverse functions that are either former components of the contractile ring and central spindle, or specifically recruited during the midbody maturation process that ultimately leads to the final physical separation or abscission of the two daughter cells (D’Avino and Capalbo, 2016; Mierzwa and Gerlich, 2014). Midbody proteins are arranged in a very precise and stereotyped spatial pattern along the midbody (Hu et al., 2012), which can be divided in approximately three major regions: the midbody ring, the midbody central core, and the midbody arms, which flank the midbody core (D’Avino and Capalbo, 2016). The proper localization, regulation and interactions of midbody proteins are essential for the execution of abscission and for preventing incorrect genome segregation (Mierzwa and Gerlich, 2014). The large multifunctional kinase CIT-K has been shown to play a key evolutionarily conserved role in maintaining the orderly arrangement of midbody proteins (D’Avino, 2017). CIT-K exerts this role by interacting and regulating the localization of several midbody proteins, including anillin, the centralspindlin complex (an heterotetramer composed of two subunits of the kinesin KIF23/MKLP1 and two molecules of RacGAP1), the chromosomal passenger complex (CPC, of which AURKB is the kinase subunit), the kinesins KIF14 and KIF20A, and the GTPase RhoA (Bassi et al., 2013; Bassi et al., 2011; Capalbo et al., 2019; Gai et al., 2011; Gruneberg et al., 2006; McKenzie et al., 2016).

Recent studies have revealed that the midbody has also important functions post-mitosis. After abscission, the midbody remnants (MBRs) can be either reabsorbed by one of the daughter cells or released into the extracellular environment and then eventually internalized by another cell (Crowell et al., 2014; Peterman et al., 2019). These post-mitotic MBRs have been implicated in diverse biological processes, including cell fate, pluripotency, apical-basal polarity, tissue organization, cell proliferation, and cilium and lumen formation (Dionne et al., 2015; Peterman and Prekeris, 2019). Recent studies have indicated that MBRs are large extracellular vesicles that contain ribonucleoprotein (RNP) complexes characterized by distinct populations of mRNAs and spatiotemporally regulated translation (Park et al., 2023; Rai et al., 2021; Suwakulsiri et al., 2024). Therefore, MBRs can function as intercellular signaling organelles that can influence the function, activity and properties of neighboring cells (Kuriyama et al., 2025). In support of this, MBRs isolated from cancer cells can promote tumorigenic properties in non-transformed cells (Rai et al., 2021). Finally, some midbody proteins have been linked to brain development and microcephaly (Siskos et al., 2021). Despite the evidence of the involvement of the midbody in these important processes, our understanding of the mechanisms that regulate its formation, functions and inheritance is very limited. Importantly, initial studies have revealed that phosphorylation plays a key role in regulating the activity, function, and associations of midbody proteins in both time and space (D’Avino and Capalbo, 2016). For example, the activity and interactions of the centralspindlin complex are regulated by complex changes in its phosphorylation status mediated by CDK1, AURKB, Plk1, and PP1 (Adriaans et al., 2019; Basant et al., 2015; Burkard et al., 2009; Capalbo et al., 2019; Douglas et al., 2010; Mishima et al., 2004; Wolfe et al., 2009). In addition, we have previously described a cross-regulation mechanism between AURKB and CIT-K that controls the assembly of the midbody by temporally regulating the association of CIT-K with both the MKLP1/KIF23 centralspindlin component and the CPC (McKenzie et al., 2016). Together, these studies highlighted that the assembly and functions of this important organelle are regulated by complex and interconnected phosphorylation events that have yet to be fully understood.

Here, we report a novel cross-regulatory mechanism among mitotic kinases important for the formation and stability of the midbody. We found that CDK1 and AURKB coordinately regulate CIT-K distribution and association with its partners by phosphorylating two distinct residues located adjacent to or within the coiled coil domain of CIT-K. Analyses using phospho-specific antibodies and phospho-mimetic and non-phosphorylatable mutants indicated that perturbing these phosphorylation events leads to abnormal midbody formation, accumulation of post-mitotic midbody remnants, and affects the interaction of CIT-K with its midbody partners. Together, our findings indicate that coordinated regulation of CIT-K by CDK1 and AURKB temporally regulates the association of CIT-K with its partners AURKB, centralspindlin, and KIF14, in order to finely dictate midbody assembly, stability and inheritance.

## Results

### CIT-K is phosphorylated by CDK1 and AURKB at multiple residues

During our previous proteomics studies of the CIT-K protein-protein interaction networks (interactomes) during mitosis (Capalbo et al., 2019; McKenzie et al., 2016) we identified several phosphorylated serine (S) and threonine (T) residues that matched the consensus sequences for CDK1 and AURKB (Fig. 1A). We previously confirmed that S699 and a stretch of two S and one T residues at positions 1385–1387 were phosphorylated by AURKB *in vitro* (McKenzie et al., 2016). We found that the fragments containing potential CDK1 phosphorylation sites, encompassing the kinase domain (CIT-K_1-423_; S400), the first coiled coil region, CC1 (CIT-K_420-785_, S440), and the C-terminal region (CIT-K_1344-2060_, S1982), were phosphorylated *in vitro* by CDK1 (Fig. 1B). The CC1 domain has been shown to interact with the kinesins KIF14 and MKLP1/KIF23 (Bassi et al., 2013; Gruneberg et al., 2006; Watanabe et al., 2013) and we previously reported that phosphorylation of S699 by AURKB regulates CIT-K localization and its association with KIF23 and the CPC (McKenzie et al., 2016). Therefore, we decided to further investigate the role of S440 phosphorylation in detail, also in relation to AURKB-mediated phosphorylation of S699. First, we confirmed that S440 is a direct CDK1 target residue because a fragment containing an S to A mutation at this residue (S440A) was no longer phosphorylated by CDK1 *in vitro* (Fig. 1C). Both S440 and S699 are conserved in vertebrates (Fig. 1D). We generated a predicted structure of the longest and only CIT-K isoform that contains S699 (UniProt ID O14578-4) using AlphaFold2 (Jumper et al., 2021). This predicted structure indicated that CIT-K forms a folded protein with the N-terminal kinase domain positioned very close to the C-terminal PH and CNH domains (Fig. 1E). Interestingly, S440 and S699 are exposed and easily accessible, and located in two structurally separated regions (Fig. 1E). In particular, S440 lies in a predicted disordered region just upstream of the CC1 domain, while S699 is located within the CC1 (Fig. 1A and E).

**Figure 1.**
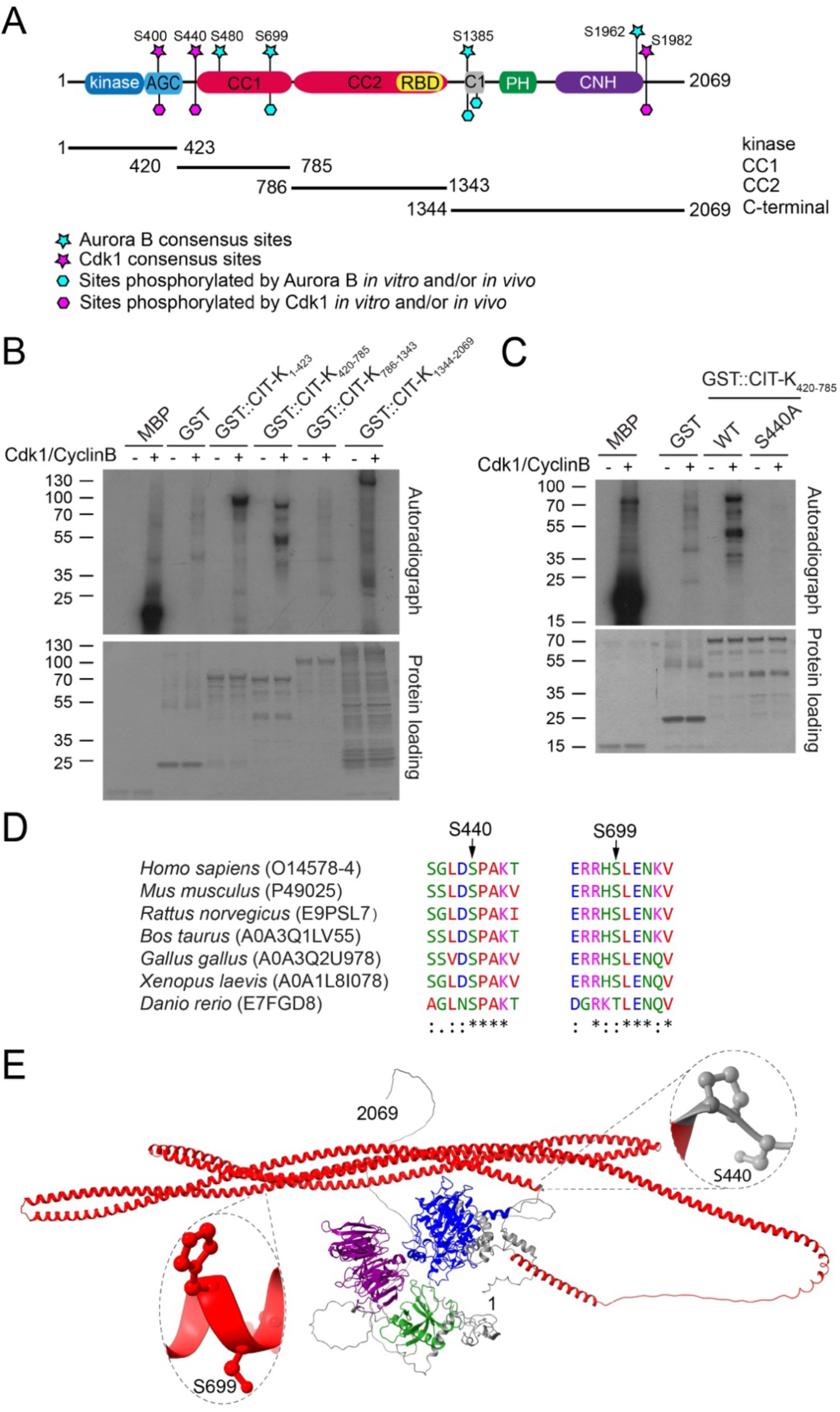
CDK1 and AURKB phosphorylate distinct CIT-K residues. (A) Schematic diagram illustrating the protein domains of CIT-K. The positions of the different CIT-K fragments used for the *in vitro* phosphorylation assays are also indicated. CC1 and CC2 indicate the fragments encompassing the first and second coiled coil regions; C1, cysteine-rich motif; PH, Pleckstrin Homology domain; CNH, Citron-Nik1 homology domain. (B) GST-tagged CIT-K fragments, GST alone or the positive control myelin basic protein (MBP) were incubated with (+) or without (−) recombinant CDK1–cyclin B complexes, in the presence of [γ-32P] ATP. The reactions were separated by SDS-PAGE, and the gel stained with Coomassie Blue to check the protein loading (shown at the bottom). The gel was then transferred onto nitrocellulose membrane and exposed to X-ray films. The numbers on the left indicate the sizes in kDa of the molecular mass marker. (C) GST-tagged wild type CIT-K_420-785_ fragment, its non-phosphorylatable mutant variant S440A, GST alone or the positive control myelin basic protein (MBP) were incubated with (+) or without (−) recombinant CDK1–cyclin B complexes, in the presence of [γ-32P] ATP. The reactions were separated by SDS-PAGE, and the gel stained with Coomassie Blue to check the protein loading (shown at the bottom). The gel was then transferred onto nitrocellulose membrane and exposed to X-ray films. The numbers on the left indicate the sizes in kDa of the molecular mass marker. (D) The amino acid sequences containing the S440 and S699 residues of human CIT-K and its orthologues in other vertebrate species were aligned using the Muscle algorithm (Edgar, 2004). Amino acids are colored according to their chemical properties. (E) Schematic diagram illustrating the CIT-K structure predicted by AlphaFold2. The kinase, CC, PH and CNH domain are colored as in (A). The position of S440 and S699 are indicated and magnified.

These results indicate that CIT-K is phosphorylated by CDK1 and AURKB at two distinct and evolutionarily conserved serine residues located adjacent or within the first coiled coil domain, which mediates the interaction of CIT-K with KIF14 and KIF23/MKLP1 (Bassi et al., 2013; Gruneberg et al., 2006; Watanabe et al., 2013).

### S440 and S699 phosphorylations display distinct temporal profiles

To analyze the temporal and spatial profiles of S440 and S699 phosphorylation, we generated antibodies directed against peptides phosphorylated at either S440 (pS440) or S699 (pS699). These antibodies specifically recognized the CIT-K_420-785_ fragment phosphorylated *in vitro* by CDK1 and AURKB, and not if the S440 and S669 residues were mutated to alanine (Fig 2A and 2B). We then analyzed the phosphorylation profiles of pS440 and pS699 by Western blotting analyses using extracts from cells synchronized at different mitotic stages through thymidine/nocodazole block and release. To assess the specificity of the two phospho-antibodies, we employed a HeLa cell line expressing a CIT-K::GFP transgene (Capalbo et al., 2019; McKenzie and D’Avino, 2016) and extracted proteins from cells depleted of endogenous CIT-K by using an siRNA directed against the 3’UTR of CIT-K, which is not present in the CIT-K::GFP transgene. Western blot analysis showed that both phospho antibodies recognized two bands in control cells and that the faster migrating band, corresponding to endogenous CIT-K, was absent in extracts from cells treated with the 3’UTR CIT-K siRNA (Fig 2C and 2E). These results indicated that the antibodies specifically recognize CIT-K phosphorylated at S440 and S699. Our results indicated that the CIT-K pS440 signal decreased 90 minutes after exit from mitosis when compared to total CIT-K, in parallel to cyclin B degradation (Fig. 2C). In addition, CIT-K pS440 was markedly reduced after treatment with the CDK1 inhibitor RO-3306 (Fig. 2D), confirming that pS440 is phosphorylated by CDK1 in cells. By contrast, pS699 showed a less marked decline compared to total CIT-K, being still present 120 minutes after release from nocodazole (Fig. 2E). CIT-K pS699 levels reduced after treatment with the AURKB inhibitor ZM447439 (Fig. 2F), further confirming that S699 is phosphorylated by AURKB in cells. Immunofluorescence analysis indicated that CIT-K pS440 was found diffuse in the cytoplasm in metaphase and anaphase, accumulated at the cleavage furrow in early telophase, and then, after completion of furrow ingression, it localized to the midbody core and arms instead of the normal localization of CIT-K to midbody ring (Fig. 3) (Bassi et al., 2013; Capalbo et al., 2019; D’Avino, 2017). Notably, the pS440 signal to the midbody core and arms did not colocalize with CIT-K::GFP and did not always disappear after CIT-K RNAi (Fig. 3), suggesting that it might not be specific for CIT-K. However, we cannot exclude that this signal may represent a relatively small, but particularly stable population of CIT-K pS440. Unfortunately, the CIT-K pS669 antibody failed to detect any specific signal that was reduced or eliminated after CIT-K RNAi, indicating that this phospho-antibody is not suitable for immunofluorescence experiments (Supplementary Fig. S1). Our findings indicate that CIT-K is phosphorylated at S440 and S699 in early mitotic stages, but while S440 is de-phosphorylated after anaphase onset, S699 remains phosphorylated also in anaphase and telophase. These results are consistent with the evidence that CDK1 activity declines after cyclin B degradation in anaphase, while AURKB remains active in cytokinesis (D’Avino and Capalbo, 2015; D’Avino and Capalbo, 2016; Lindon et al., 2015; Wieser and Pines, 2015).

**Figure 2.**
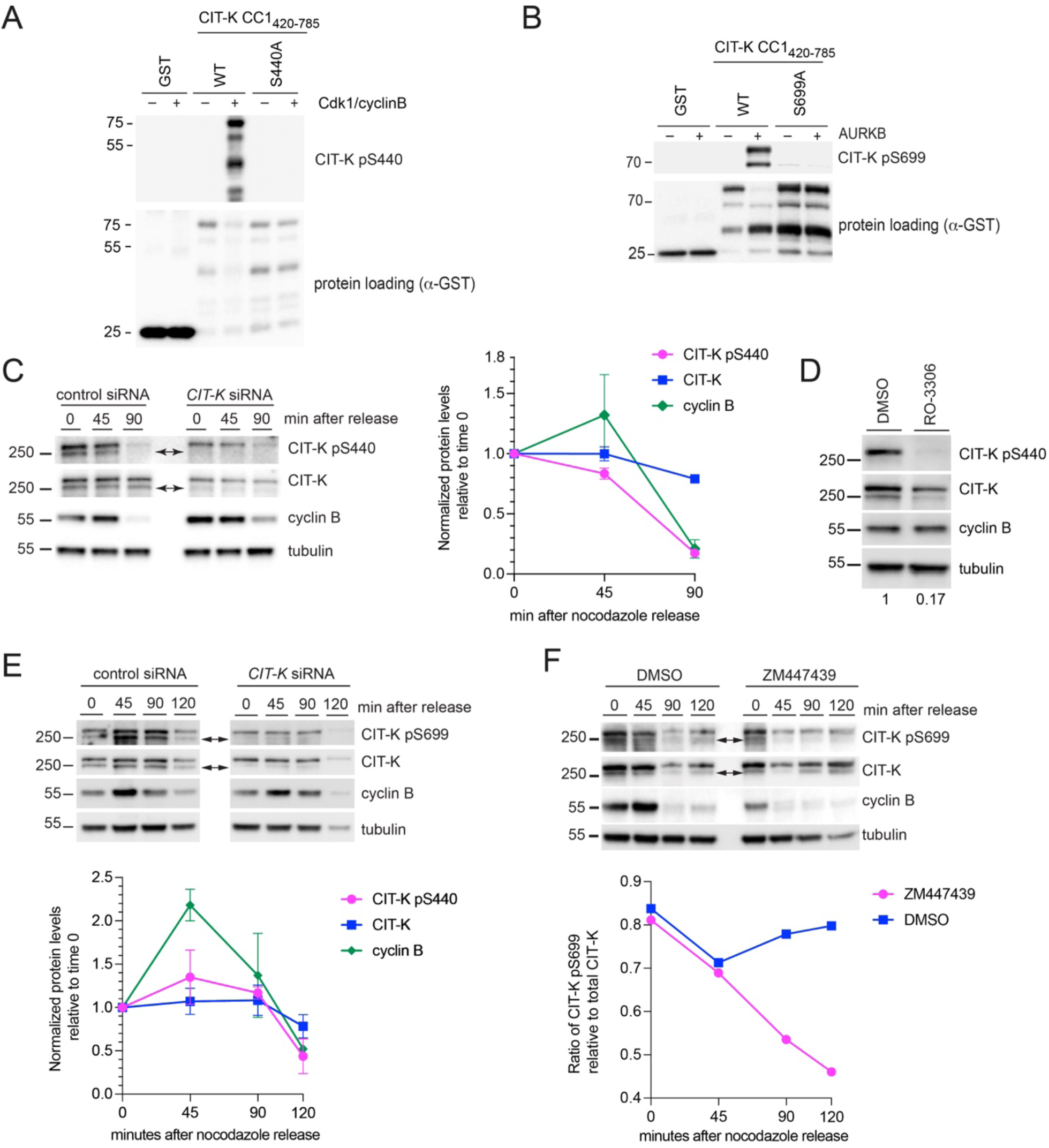
Temporal profiles of S440 and S699 phosphorylation. (A) GST-tagged wild type CIT-K_420-785_ fragment, its non-phosphorylatable mutant variant S440A, and GST alone were incubated with (+) or without (-) recombinant CDK1/cyclin B. The reactions were then separated by SDS PAGE and analyzed by Western Blot to detect phosphorylated S440 (pS440) and GST. The numbers on the left indicate the sizes in kDa of the molecular mass marker. (B) GST-tagged wild type CIT-K_420-785_ fragment, its non-phosphorylatable mutant variant S699A, and GST alone were incubated with (+) or without (-) recombinant Aurora B (AURKB). The reactions were then separated by SDS PAGE and analyzed by Western Blot to detect phosphorylated S699 (pS699) and GST. The numbers on the left indicate the sizes in kDa of the molecular mass marker. (C) HeLa Kyoto cells stably expressing GFP-tagged CIT-K were treated with siRNAs directed against either a random sequence (control) or the 3’UTR of *CIT-K*. During RNAi incubation, cells were synchronized by thymidine/nocodazole block and then collected at the indicated time points after nocodazole release. Proteins were extracted and analyzed by Western blot to detect CIT-K, CIT-K pS440, cyclin B and tubulin as loading control. The arrows indicated endogenous untagged CIT-K. The numbers on the left indicate the sizes in kDa of the molecular mass marker. The graph on the right shows the quantification of protein levels, normalized to tubulin and relative to the 0 time point, from at least two different Western blots. (D) HeLa Kyoto cells stably expressing GFP-tagged CIT-K were treated with siRNAs directed against either a random sequence (control) or the 3’UTR of *CIT-K*. During RNAi incubation, cells were synchronized by thymidine/nocodazole block, released for 45 min and incubated for further 30 min with either the CDK1 inhibitor RO-3306 or its solvent DMSO as control. Proteins were extracted and analyzed by Western blot to detect CIT-K, CIT-K pS440, cyclin B and tubulin as loading control. The numbers at the bottom indicated the quantification of the normalized levels of CIT-K pS440 relative to CIT-K. The numbers on the left indicate the sizes in kDa of the molecular mass marker. (E) HeLa Kyoto cells stably expressing GFP-tagged CIT-K were treated with siRNAs directed against either a random sequence (control) or the 3’UTR of *CIT-K*. During RNAi incubation, cells were synchronized by thymidine/nocodazole block and then collected at the indicated time points after nocodazole release. Proteins were extracted and analyzed by Western blot to detect CIT-K, CIT-K pS699, cyclin B and tubulin as loading control. The arrows indicated endogenous untagged CIT-K. The numbers on the left indicate the sizes in kDa of the molecular mass marker. The graph below shows the quantification of protein levels, normalized to tubulin and relative to the 0 time point, from at least two different Western blots. (F) HeLa Kyoto cells stably expressing GFP-tagged CIT-K were treated with siRNAs directed against either a random sequence (control) or the 3’UTR of *CIT-K*. During RNAi incubation, cells were synchronized by thymidine/nocodazole block, released and incubated with either the AURKB inhibitor ZM447439 or its solvent DMSO. Cells were collected at the indicated time points and proteins were extracted and analyzed by Western blot to detect CIT-K, CIT-K pS699, cyclin B and tubulin as loading control. The arrows indicated endogenous untagged CIT-K. The numbers on the left indicate the sizes in kDa of the molecular mass marker. The graph below shows the quantification of protein levels, normalized to tubulin and relative to the 0 time point, of the ratio of CIT-K pS699 versus endogenous CIT-K.

**Figure 3.**
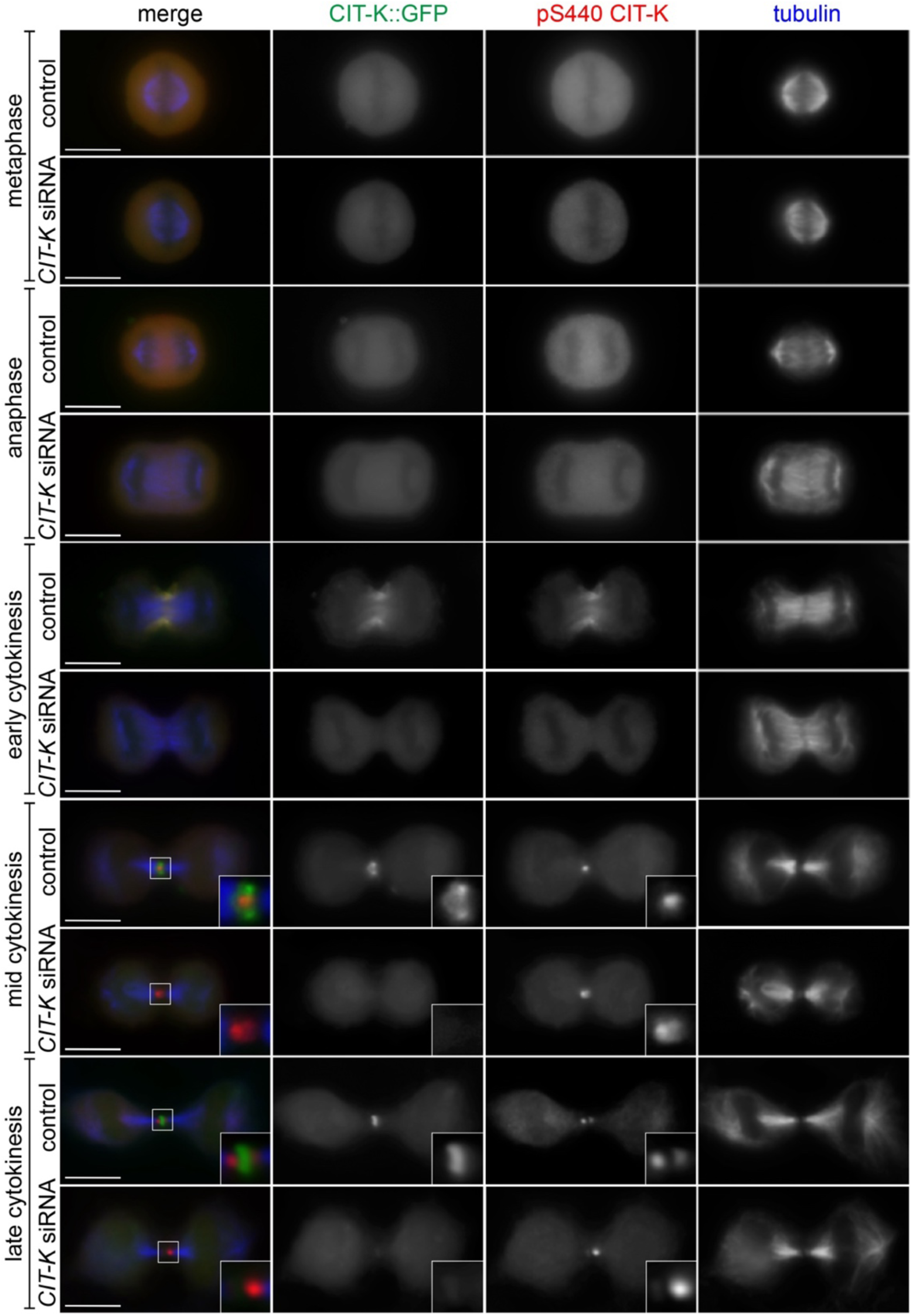
Localization of CIT-K phosphorylated at S440 during cell division. HeLa Kyoto cells stably expressing GFP-tagged CIT-K were treated with siRNAs directed against either a random sequence (control) or the 3’UTR of *CIT-K* and after 48 hours were fixed and stained to detect CIT-K::GFP (green), CIT-K pS440 (red) and tubulin (blue). The shape and thickness of microtubule bundles at the intercellular bridge were used as criteria to stage telophase cells. Insets show a 3X magnification of the midbody. Bars, 10 µm.

### Phosphorylation at S440 regulates midbody formation and stability

To investigate the role of S440 and S699 phosphorylation during cell division, we generated HeLa cell lines stably expressing doxycycline-inducible GFP-tagged CIT-K transgenes carrying wild type (WT), phosphomimetic (S∏D), or phospho-dead (S∏A) mutations at these residues. We selected individual clones that expressed the transgenes at levels that were most similar to that of endogenous CIT-K after testing different concentrations of doxycycline (Supplementary Fig. S2). We analyzed the distribution of these mutant transgenes in both control cells and cells depleted of the endogenous CIT-K using an siRNA targeting the 3’UTR region of the gene, which is not present in the transgenes. Both mutant proteins localized to the midbody in late cytokinesis (Supplementary Figure S3). However, S440 mutant CIT-K proteins often failed to assemble in the ring-like structure typical of CIT-K and showed excessive accumulation to the midbody in very late cytokinesis, just before abscission, a phenotype that was more penetrant for the S440D mutant (Supplementary Fig. S3B-C). We then tested the ability of these mutants to rescue the multinucleation (a readout of cytokinesis failure) caused by RNAi depletion of endogenous CIT-K and found that only the S699A mutant failed to rescue (Fig. 4A). We also determined the frequency of daughter cells linked by an intercellular bridge (cells in cytokinesis), as a readout for possible abscission delays. Depletion of CIT-K in the parental cells led to a decrease of cells in cytokinesis (Fig. 4B), which is consistent with the established role of CIT-K in midbody formation (Bassi et al., 2013; McKenzie et al., 2016). Both S440A and S440D mutants showed a significant increase of cells in cytokinesis, which in the case of S440D occurred also in the presence of endogenous CIT-K, indicating a dominant effect (Fig. 4B). More modest increases in cell in cytokinesis were found with S669A and S669D mutants after 3’UTR CIT-K siRNA depletion, but they were not significant when compared to the WT transgene (Fig. 4B). We also found that the distribution of S440A and S440D mutant proteins to the midbody appeared disorganized and fragmented in late cytokinesis and post-abscission (Fig. 4C and 4D). Co-staining with the midbody components and CIT-K interactors AURKB, KIF14 and MKLP1/KIF23 confirmed that these represented entire midbodies and not just aggregates of the GFP::CIT-K mutant proteins. Notably, the midbody distribution of these three CIT-K interactors was altered in cells expressing S440A and S440D mutants (Fig. 4C). In particular, AURKB appeared to unusually persist at the midbody after abscission in both cell lines and its signal was also abnormally observed at the midbody core where it partially colocalized with CIT-K mutant proteins (Fig. 4C). These incorrectly assembled midbodies were more frequent in cells expressing the S440A mutant, but more severe in cells expressing S440D (Fig. 4C and 4D). Finally, we found that interphase cells expressing the S440A and S440D CIT-K mutant proteins contained more midbody remnants (MBRs) (Fig. 4E and 4F). Midbody fragmentation and increased MBRs were not observed in cells expressing S699A or S699D mutants (Fig. 4D and 4F).

**Figure 4.**
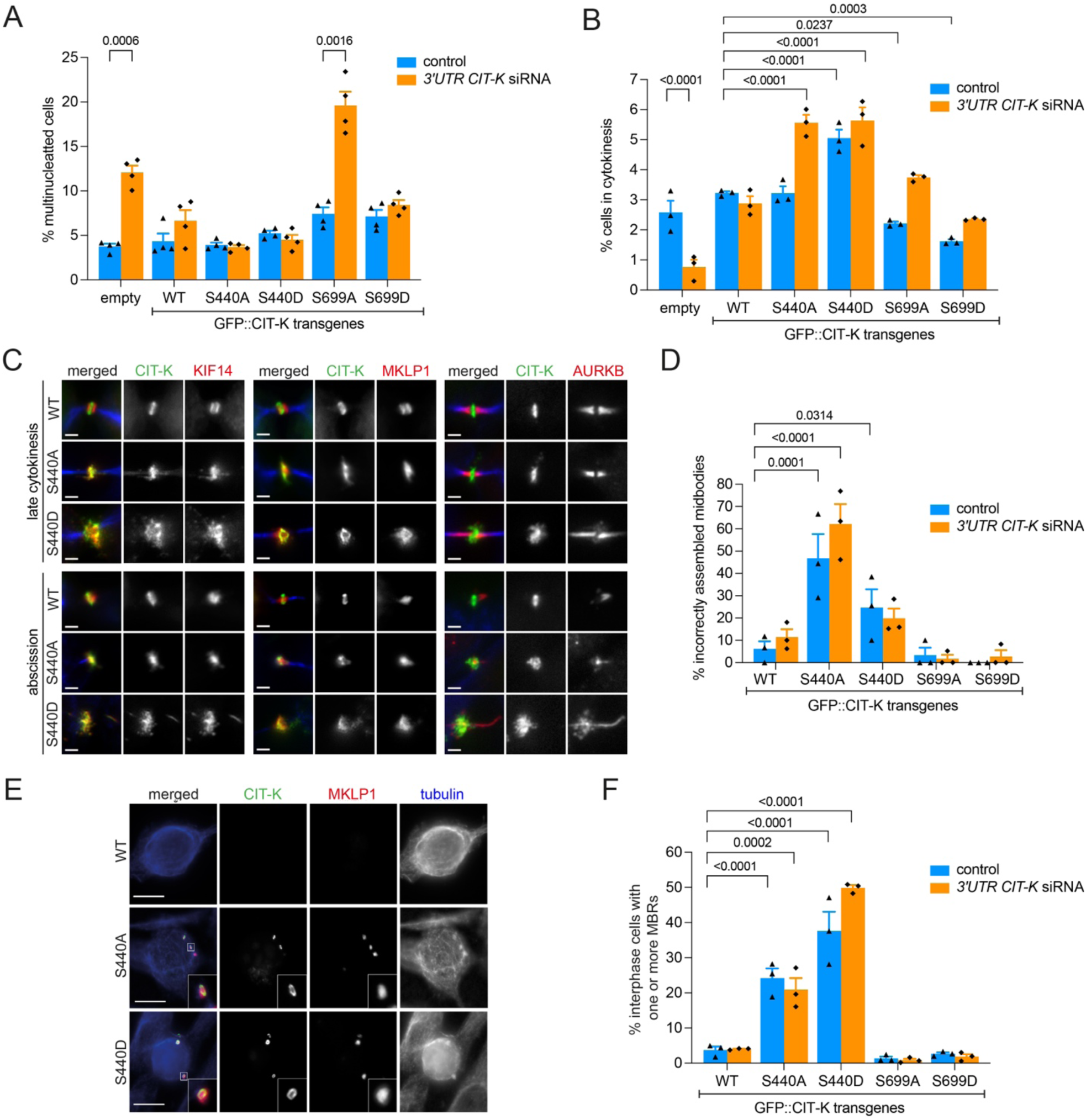
CIT-K phosphorylation at S440 and S699 regulates midbody structure and stability. (A-B) HeLa cells expressing the indicated GFP::CIT-K constructs or no transgene (empty) were transfected with either an siRNA targeting the 3’UTR of *CIT-K* with siRNAs directed against either a random sequence (control) or the 3’UTR of *CIT-K* and after 48 hours were fixed and stained to detect tubulin and DNA. The samples were then analyzed to quantify the percentage of multinucleated cells (A) and daughter cells linked by an intercellular bridge (cells in cytokinesis) (B). More than 1000 cells were counted in n≥3 independent experiments. Bars indicate standard errors. Significant *p* values are indicated at the top of the graph (unpaired t-test in A and two-way ANOVA in B). (C) HeLa cells expressing the indicated GFP::CIT-K constructs were treated with siRNA directed against the 3’UTR of *CIT-K* and after 48 hours were fixed and stained to detect CIT-K (green), tubulin (blue) and KIF14, MKLP1 or AURKB (red). The shape and thickness of microtubule bundles at the intercellular bridge were used as criteria to stage cells. Bars, 2 µm. (D) Quantification of late cytokinesis cells showing disorganized and/or fragmented midbodies from the experiments shown in A-C. More than 500 cells were counted in n≥3 independent experiments. Bars indicate standard errors. Significant *p* values are indicated at the top of the graph (two-way ANOVA). (E) HeLa cells expressing the indicated GFP::CIT-K constructs were treated with siRNA directed against the 3’UTR of *CIT-K* and after 48 hours were fixed and stained to detect CIT-K (green), tubulin (blue) and MKLP1 (red) and then analyzed to quantify the number of midbody remnants. Bars, 10 µm. (F) Quantification of interphase cells from the experiments shown in E containing one or more midbody remnants (MBRs). More than 500 cells were counted in n≥3 independent experiments. Bars indicate standard errors. Significant *p* values are indicated at the top of the graph (two-way ANOVA).

Together, these results indicate that S440 phosphorylation has a more relevant role on midbody formation and stability, and in turn on abscission, than phosphorylation at S699, which instead seems to be more important for CIT-K localization and successful cytokinesis.

### Phosphorylation at S440 and S699 regulates CIT-K association with its midbody partners

To gain insights into the possible underlying molecular mechanisms of CIT-K regulation via S440 and S699 phosphorylation, we investigated the ability of the S440 and S699 phospho-dead and phospho-mimetic mutants to interact with known CIT-K midbody partners, AURKB, KIF14, and MKLP1/KIF23, by immunoprecipitation using proteins extracted from cells synchronized in telophase and depleted of endogenous CIT-K. The S440A phospho-dead mutation slightly increased the association of CIT-K::GFP with its midbody partners (Fig. 5A and B). By contrast, the S440D mutation considerably reduced the ability of CIT-K to pull down any of its partners (Fig. 5A and 5B). In agreement with our previous findings (McKenzie et al., 2016), the S699A mutant pulled down all three CIT-K midbody partners much more efficiently than the WT counterpart (Fig. 5C and 5D). The S699D mutant pulled down the three partners less efficiently than S699A, but more strongly than the WT (Fig. 5C and 5D).

**Figure 5.**
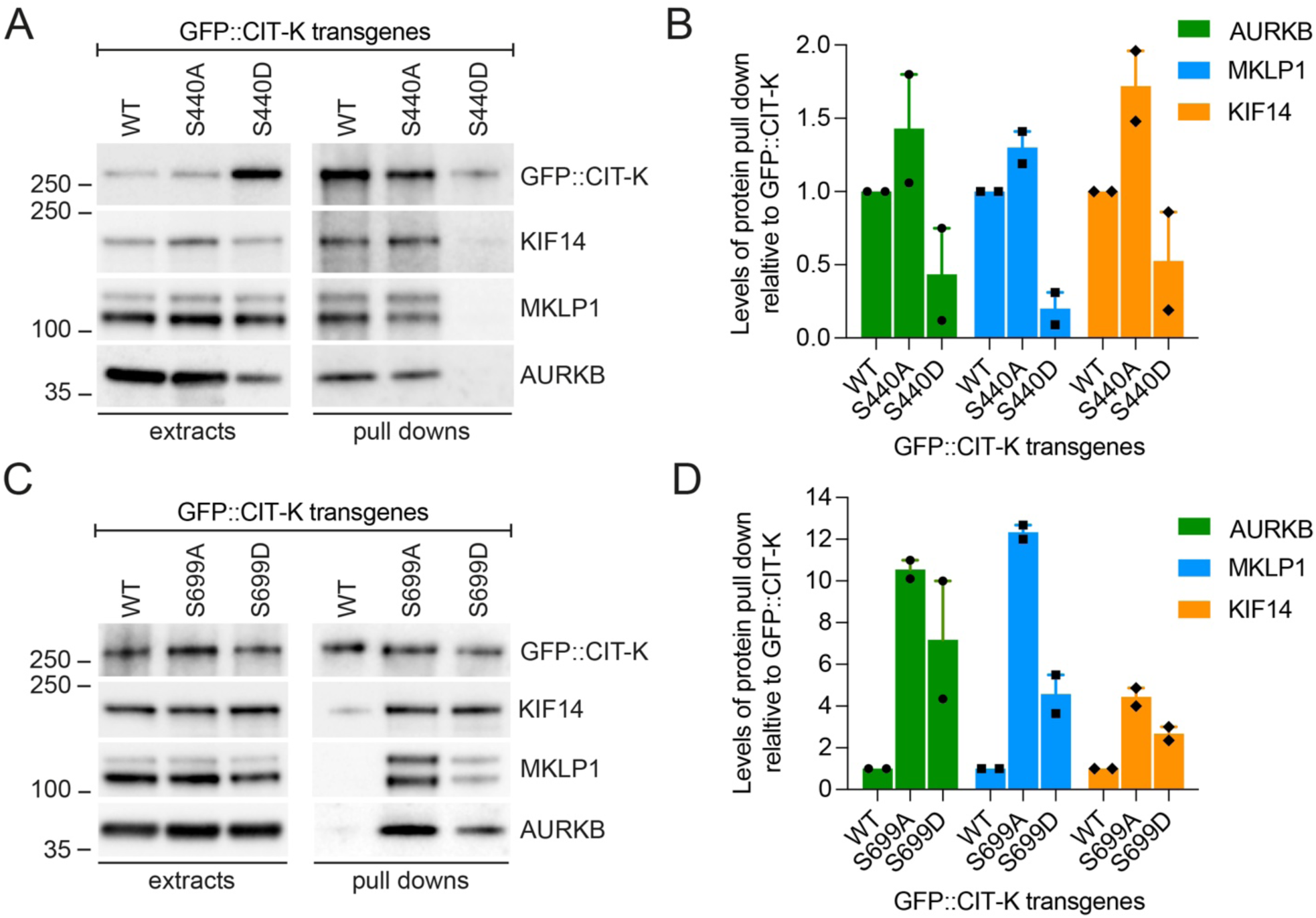
Phosphorylation at S440 and S699 affects CIT-K interaction with its midbody partners. (A) HeLa Kyoto cells stably expressing the indicated GFP-tagged CIT-K transgenes were treated with an siRNA directed against the 3’UTR of *CIT-K*. During RNAi incubation, cells were synchronized by thymidine/nocodazole block and then collected 90 minutes after nocodazole release. Protein extracts were used in a GFP pull-down assay and then analyzed by Western blot to detect the proteins indicated at the right. The numbers on the left indicate the sizes in kDa of the molecular mass marker. (B) Quantification of protein levels from experiments as shown in A, normalized to tubulin and relative to GFP::CIT-K transgenes, from two different Western blots. Bars indicate standard errors. (C) HeLa Kyoto cells stably expressing the indicated GFP-tagged CIT-K transgenes were treated with an siRNA directed against the 3’UTR of *CIT-K*. During RNAi incubation, cells were synchronized by thymidine/nocodazole block and then collected 90 minutes after nocodazole release. Protein extracts were used in a GFP pull-down assay and then analyzed by Western blot to detect the proteins indicated at the right. The numbers on the left indicate the sizes in kDa of the molecular mass marker. (D) Quantification of protein levels from experiments as shown in C, normalized to tubulin and relative to GFP::CIT-K transgenes, from two different Western blots. Bars indicate standard errors.

CIT-K directly binds to KIF14 and MKLP1 through its CC1 domain (Bassi et al., 2013; McKenzie et al., 2016; Watanabe et al., 2013), whereas AURKB and other CPC components binds to the CNH domain (McKenzie et al., 2016). As both S440 and S699 are located near or within the CC1 region, we investigated whether the ability of CIT-K to bind to KIF14 and/or MKLP1/KIF23 *in vitro* was affected by phosphorylation at these two residues. We divided the fragment containing the CC1 domain, CIT-K_420-785_, into two smaller peptides, CIT-K_420-585_ and CIT-K_586-785_, the first containing the S440 residue and the other S699 (Fig. 6A). These peptides were tagged with the maltose-binding protein (MBP), expressed and purified from bacteria, and then tested in an *in vitro* pull down assay for their ability to bind to bacterially-purified GST-tagged KIF14_901-1648_ and MKLP1_620-856_ peptides, known to directly interact with CIT-K (Bassi et al., 2013; Watanabe et al., 2013). GST:: KIF14_901-1648_ specifically pulled down the CIT-K_420-585_ fragment containing S440 (Fig. 6B), while GST:: MKLP1_620-856_ specifically pulled down the other peptide, CIT-K_586-785,_ containing S699 (Fig. 6C). We then tested whether phosphorylation at S440 or S699 affected the binding of these CIT-K fragments to KIF14 or MKLP1. S440 phosphorylation did not appear to affect the binding of CIT-K_420-585_ to KIF14 (Fig. 6D), while S699 phosphorylation consistently reduced the association of CIT-K_586-785_ to MKLP1 *in vitro* (Fig. 6E).

**Figure 6.**
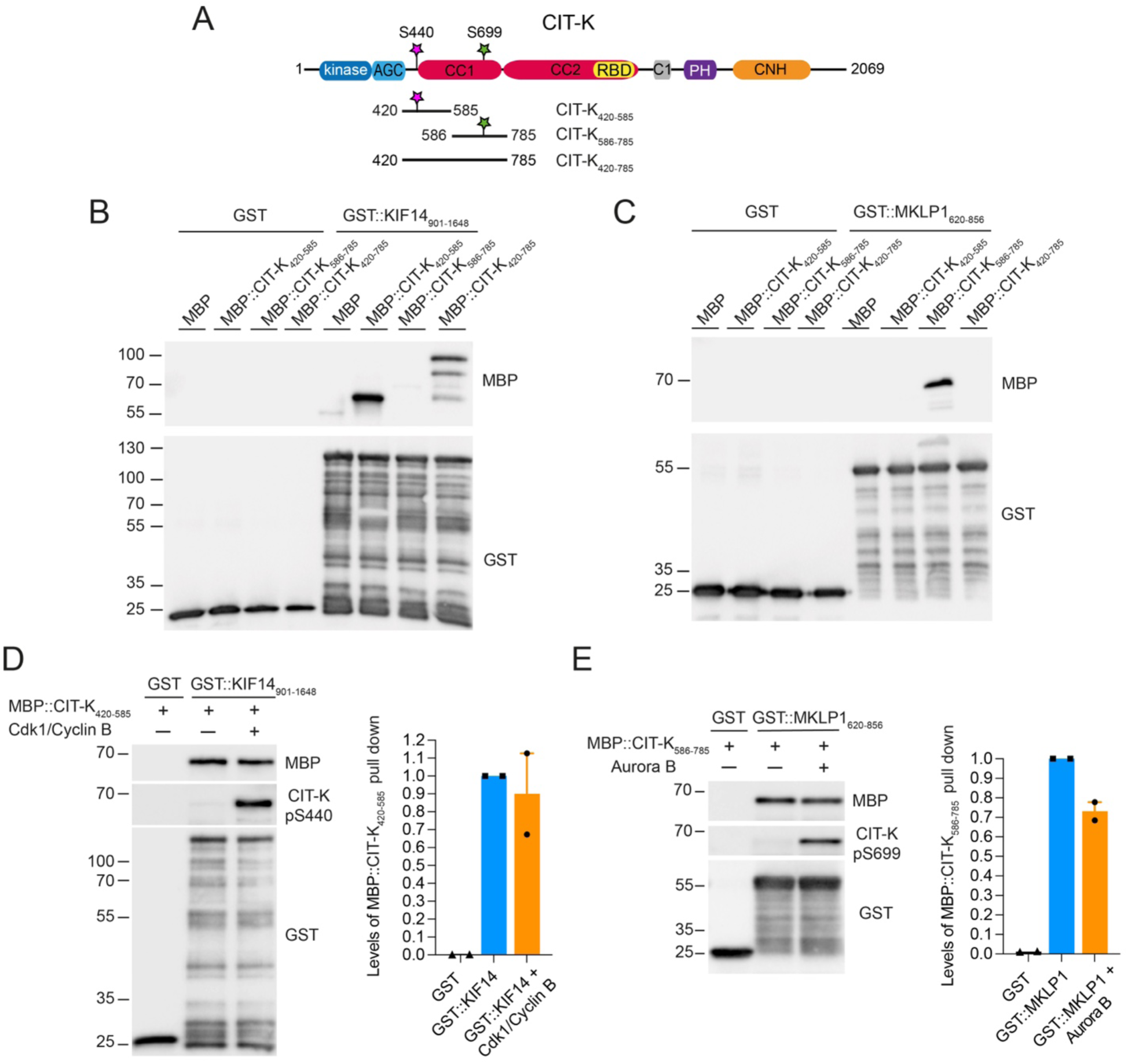
KIF14 and KIF23/MKLP1 bind to distinct regions within the first coiled coil domain of CIT-K. (A) Schematic diagram illustrating the protein domains of CIT-K. The S440 and S699 residues and the different CIT-K fragments used for the *in vitro* pull down assays are indicated. CC1 and CC2 indicate the fragments encompassing the first and second coiled coil regions; C1, cysteine-rich motif; PH, Pleckstrin Homology domain; CNH, Citron-Nik1 homology domain. (B) The GST::KIF14_901-648_ fragment and GST alone were incubated with the indicated MBP-tagged CIT-K fragments, and then pulled down using glutathione beads. The pull downs were then analyzed by Western blot. The numbers on the left indicate the sizes in kDa of the molecular mass markers. (C) The GST::MKLP1_620-856_ fragment and GST alone were incubated with the indicated MBP-tagged CIT-K fragments, and then pulled down using glutathione beads. The pull downs were then analyzed by Western blot. The numbers on the left indicate the sizes in kDa of the molecular mass markers. (D) The MBP::CIT-K_420-585_ fragment was incubated with (+) or without (−) recombinant CDK1-cyclin B and then used in pull down assays with the GST::KIF14_901-648_ fragment and GST alone. The pull downs were analyzed by Western blot using antibodies against MBP, CIT-K pS440, and GST. The numbers on the left indicate the sizes in kDa of the molecular mass markers. The graph on the right shows the quantification of the protein levels of MBP::CIT-K_420-585_, normalized to GST::KIF14_901-648_ or GST alone, and relative to unphosphorylated (-) MBP::CIT-K_420-585_. Bars indicate standard errors. (D) The MBP::CIT-K_586-785_ fragment was incubated with (+) or without (−) recombinant Aurora B and then used in pull down assays with the GST::MKLP1_620-856_ fragment and GST alone. The pull downs were then analyzed by Western blot using antibodies against MBP, CIT-K pS699, and GST. The numbers on the left indicate the sizes in kDa of the molecular mass markers. The graph on the right shows the quantification of the protein levels of MBP::CIT-K_586-785_, normalized to GST::MKLP1_620-856_ or GST alone, and relative to unphosphorylated (-) MBP::CIT-K_586-785._ Bars indicate standard errors.

Taken together, our findings indicate the phosphorylation at S440 and S699 reduces the ability of CIT-K to interact with its midbody partners AURKB, KIF14 and KIF23/MKLP1 in cells. *In vitro* binding assays indicate that S440 phosphorylation does not affect the binding of CIT-K to KIF14, while S699 phosphorylation causes a reduction of the binding of CIT-K to MKLP1.

## Discussion

CIT-K is a one of the key factors that control midbody assembly and architecture in late cytokinesis (D’Avino, 2017). We previously showed that CIT-K is necessary to maintain the orderly arrangement of midbody proteins and that cross-regulation between CIT-K and AURKB regulates midbody formation (Bassi et al., 2013; McKenzie et al., 2016). More specifically, AURKB phosphorylation of CIT-K at S699 prevents its accumulation at the spindle midzone in early cytokinesis by inhibiting the association of CIT-K with KIF23/MKLP1 and the CPC (McKenzie et al., 2016). Here, we expand these previous findings by showing that CIT-K is also phosphorylated by CDK1 at S440 and that phosphorylation by these two kinases coordinately regulates CIT-K distribution and its association with midbody partners (Figs 4 and 5) (McKenzie et al., 2016). The temporal profiles of S440 and S699 phosphorylation reflect the activity of the kinases responsible for their phosphorylation. Both residues are phosphorylated in early mitosis, but S440 is dephosphorylated earlier, at anaphase onset, in parallel with CDK1 inactivation (Figs 2 and 3). Our functional analyses using phospho-mimetic and non-phosphorylatable mutants indicate that these phosphorylation events have both convergent and distinct effects. They confirm and expand our previous findings that S699 phosphorylation by AURKB inhibits the association of CIT-K with some of its midbody partners (Fig.5) (McKenzie et al., 2016). This mechanism appears necessary for regulating CIT-K function and cytokinesis progression because the S699A mutant failed to rescue cytokinesis failure caused by CIT-K depletion (Fig. 4A). Results from the experiments with S440 phospho mutants indicate that CDK1 phosphorylation also inhibits the interaction of CIT-K with its midbody partners, even more efficiently than AURKB phosphorylation (Fig. 5). Together, these findings suggest that CDK1 and AURKB phosphorylations coordinately regulate CIT-K interactions to accurately set the timing of midbody formation. They may also prevent premature binding of CIT-K to KIF23/MKLP1 in early cytokinesis, which might interfere with centralspindlin clustering and central spindle assembly (Douglas et al., 2010; Hutterer et al., 2009). However, S440 mutants rescued CIT-K depletion (Fig. 4A) and did not cause incorrect localization of CIT-K to the central spindle in early cytokinesis like the S669A mutant (McKenzie et al., 2016). One possible explanation for this discrepancy could be that in early mitosis AURKB phosphorylation might be sufficient for regulating CIT-K in absence of S440 phosphorylation. To test this hypothesis, we considered the possibility of studying the effects of S440 and S699 mutant combinations, but unfortunately our attempts to generate cell lines expressing any of the possible double mutant combinations were unsuccessful. In addition, results from these experiments might be difficult to interpret.

The S440 and S699 residues are located in two distinct structural regions that specifically bind to KIF14 or KIF23/MKLP1 *in vitro* (Figs. 1 and 6). The results of these *in vitro* binding assays are only partially consistent with the outcome of the immunoprecipitation assays in cells expressing the S440 and S699 phospho mutants (Fig. 5). For example, the S440 phospho-mimetic mutant drastically reduced the association of CIT-K with KIF14, AURKB and KIF23/MKLP1 in cells (Fig. 5), but the peptide containing this residue binds only to KIF14 *in vitro* and S440 phosphorylation does not seem to affect this binding (Fig. 6). Similarly, the S699 phospho-dead mutant interacted much more efficiently with AURKB, KIF14 and KIF23/MKLP1 (Fig. 5), but the fragment harboring this serine binds specifically to KIF23/MKLP1 *in vitro* and S699 phosphorylation only slightly reduced this interaction (Fig. 6). Together, these findings indicate that phosphorylation at these two residues in cells must have a more widespread effect on CIT-K’s ability to interact with its partners than just local inhibition. For example, these phosphorylations could have knock-on effects on sequential binding events and/or on the overall CIT-K structure.

Our findings indicate that perturbing S440 phosphorylation has an impact on midbody formation and stability (Fig. 4). Cells expressing phospho-dead and phospho-mimetic S440 CIT-K mutants presented disorganized and fragmented midbodies, characterized by incorrect distribution of CIT-K and its partners, AURKB, KIF14 and KIF23/MKLP1 (Fig. 4C-D). These defects in midbody formation are accompanied by an increase in the frequency of cells in late cytokines (Fig. 4B), which most likely reflects a delay in abscission as a consequence of incorrectly assembled midbodies. Finally, the increase of MBRs in interphase cells expressing S440 phospho mutant proteins (Fig. 4E-F) indicate that MBRs are more stable and persist for longer. Notably, all these phenotypes were not observed in cells expressing S699 CIT-K mutant proteins. Together, these results indicate that S440 phosphorylation has a role in regulating CIT-K functions important for midbody assembly and MBR stability. These observations are unexpected because CDK1 activity and S440 phosphorylation decline in anaphase (Fig. 2C). It is possible to speculate that the phospho-mimetic S440D mutation might have a dominant-negative effect, because it would keep CIT-K in a pre-anaphase status, thereby sustaining inhibition of CIT-K binding to its partners and affecting proper midbody formation. Expression of the phospho-dead S440A mutant also affected midbody formation, although with less severe phenotypes (Fig. 3C). Possible explanations are that the S440A mutation does not act as expected or that inhibiting S440 phosphorylation before anaphase affects CIT-K function in cytokinesis. In addition, we cannot exclude the possibility that S440 might be phosphorylated in late cytokinesis by either CDK1 or another kinase. This would be consistent with the evidence that cyclin B2 has been shown to accumulate at the intercellular bridge in late cytokinesis (Mathieu et al., 2013), CDK1 was identified in two independent proteomic analyses of the midbody (Capalbo et al., 2019; Skop et al., 2004), and CDK1 inhibitors delay abscission (Mathieu et al., 2013). Furthermore, it might explain the detection of CIT-K pS440 persistent signals at the midbody core and arms in late cytokinesis cells (Fig. 3). However, our Western blot analyses indicate that this pool of phosphorylated S440 CIT-K at the midbody would not be very abundant (Fig. 2).

In conclusion, here we present evidence that coordinated phosphorylation of CIT-K by CDK1 and AURKB regulates midbody formation and MBR stability by dictating the timing and strength of CIT-K interaction with its partners. These findings not only expand and refine our understanding of the molecular mechanisms that regulate abscission, but can help unravelling how MBR stability is regulated, which in turn may lead to further insights into the role of MBRs in post-mitotic events and carcinogenesis.

## Material and Methods

### Molecular Biology

The clones containing different CIT-K fragments were generated as previously described (McKenzie et al., 2016). The QuikChange Lightning Site-Directed Mutagenesis Kit (Agilent) was used to generate the S440 and S699 phospho-dead (S to A) and phospho-mimetic (S to E) CIT-K mutants. For inducible expression of GFP-tagged CIT-K constructs in HeLa T-REx cells (ThermoFisher), DNA fragments were cloned into the pcDNA™FRT⁄TO vector (ThermoFisher). Sequences of all DNA constructs were verified by sequencing (Source BioScience).

### *In vitro* phosphorylation assay

DNA fragments coding for CIT-K fragments were generated by PCR and cloned into pDEST15 (ThermoFisher) to express N-terminal GST-tagged polypeptides in *E. coli*. Purified GST-tagged CIT-K fragments were incubated with 150 ng of human recombinant CDK1/Cyclin B (ThermoFisher), 0.1 mM ATP (Sigma-Aldrich), 5 μCi of [γ-^32^P] ATP (6000 Ci/mmol, 10 mCi/ml) (PerkinElmer) and kinase buffer (20 mM HEPES pH 7.5, 2 mM MgCl_2_, 1 mM DTT) in a final reaction volume of 15 μl. After 30 min incubation at 30°C with constant agitation, 15 μl of 2x Laemmli sample buffer was added to stop the reaction. Samples were boiled for 10 min and loaded on a 4-20% Tris-Glycine precast gel (ThermoFisher). Gels were stained with Quick Coomassie Stain (Generon) to check the protein loading and then proteins were transferred onto a nitrocellulose membrane using the iBlot Dry Blotting System (Invitrogen). Membranes were exposed to Kodak BioMax XAR Films (Sigma) at −80°C.

Non-radioactive *in vitro* kinase assays were performed as above, but without [γ-^32^P] ATP and by using 150 ng of human recombinant CDK1/Cyclin B or 190 ng of human recombinant Aurora B kinase (ThermoFisher) and ATP at a final concentration of 0.5 mM.

### Cell culture, siRNA transfection, drug treatments, and generation of stable cell lines

HeLa Kyoto were maintained in DMEM (Sigma) containing 10% (v/v) Fetal Bovine Serum (Sigma) and 1% (v/v) penicillin/streptomycin (Invitrogen) at 37°C and 5% CO_2_. HeLa T-REx cells (ThermoFisher) were cultured in DMEM (Sigma) containing 10 % (v/v) Fetal Bovine Serum without tetracycline (FBS-TET; Sigma) and 1 % (v/v) penicillin/streptomycin at 37°C and 5% CO_2_. HeLa T-REx transfected cell lines were maintained in the same medium containing 5 μg/ml blasticidin S and 250 μg/ml hygromycin B (ThermoFisher).

For RNA interference the following siRNAs were used: scrambled sequence control, 5’-AACGTACGCGGAATACTTCGA-3’; and *3’ UTR CIT-K*, 5’-CACACUAUGGAACUCUGCU-3, which were transfected using Lipofectamine RNAiMAX (ThermoFisher) following the manufacturer’s instructions. These siRNAs have been previously validated for specificity and efficacy (McKenzie et al., 2016).

To generate HeLa T-Rex inducible cell lines expressing WT or S440 and S699 mutant CIT-K constructs, 2 ×10^6^ cells were transfected with 10 μg of the relevant plasmid and the pOG444 plasmid in 7:3 ratio using the Neon Transfection System (ThermoFisher) and the following settings: 1005 v, 35 ms, 2 pulses. After 48 h, cells in 100 mm culture dish were selected in complete medium containing 5 μg/ml blasticidin S and 250 μg/ml hygromycin B for approximately two weeks until colonies became visible. Individual colonies were picked, cultured under resistance and tested for expression of the construct by Western blot and immunofluorescence. HeLa cells were synchronized in metaphase and telophase by using the thymidine-nocodazole block and release procedure previously described (Capalbo et al., 2019; McKenzie et al., 2016).

### Fluorescence microscopy

HeLa cells were grown on microscope glass coverslips (Menzel-Gläser) and fixed in either PHEM buffer (60 mM Pipes, 25 mM Hepes pH 7, 10 mM EGTA, 4 mM MgCl_2_, 3.7% [v/v] formaldheyde) for 12 min at room temperature or in ice-cold methanol for 10 min at −20°C. They were then washed three times for 5 min with PBS and incubated in blocking buffer (PBS, 0.5% [v/v] Triton X-100 and 5% [w/v] BSA) for 1 h at room temperature. Coverslips were incubated overnight at 4°C with primary antibodies diluted in PBT (PBS, 0.1% [v/v] Triton X-100 and 1% [w/v] BSA). The day after, coverslips were washed twice for 5 min in PBT, incubated with secondary antibodies diluted in PBT for 2h at room temperature and then washed twice with PBT and once with PBS. Coverslips were mounted on SuperFrost Microscope Slides (VWR) using VECTASHIELD Mounting Medium containing DAPI (Vector Laboratories). Phenotypes were blindly scored by at least two people independently. Images were acquired using a Zeiss Axiovert epifluorescence microscope equipped with µManager software. Fiji (Schindelin et al., 2012) was used to generate maximum intensity projections, which were adjusted for contrast and brightness, and assembled using Photoshop.

### Antibodies

CIT-K phospho-specific antibodies were raised in rabbits against synthetic peptides containing either a phosphorylated serine at position 440 (RSESVVSGLDpSPAKTSSMEKK; S440) or a phosphorylated serine at position 699 (EAEERRHpSLENKVKR, S699). Sera were first eluted through a column containing a non-phosphorylated peptide and then each antibody was affinity purified using the appropriate phospho-peptide. Peptide synthesis, conjugation, rabbit immunizations, serum production, and affinity purifications were carried out by Generon, UK.

Other antibodies used in this study and their dilutions for Western blot (WB) and immuno-fluorescence (IF) analyses were: mouse monoclonal anti α-tubulin (clone DM1A, Sigma, T9026 dilutions for WB 1:20000, for IF 1:2000), rabbit polyclonal anti-β-tubulin (Abcam, ab6046 dilutions for WB 1:5000, for IF 1:400), mouse monoclonal anti-cyclin B1 (clone GNS1, Santa Cruz, sc-245 dilution for WB 1:2000), mouse monoclonal anti-CIT-K (BD Transduction Laboratories, 611377 dilutions for WB 1:1500, for IF 1:250), rabbit polyclonal anti-KIF14 (Bethyl laboratories A300-233A dilution for WB 1:2000), rabbit polyclonal anti-MKLP1 (clone N19, Santa Cruz Biotechnology, sc-867, dilutions for WB 1:3000, for IF 1:500), rabbit recombinant monoclonal anti-MKLP1 antibody (clone EPR10879, Abcam, ab174304, dilutions for WB 1:5000, for IF 1:800), mouse monoclonal anti-Aurora B (clone AIM-1, BD Transduction Laboratories, 611082 dilutions for WB 1:2000, for IF 1:100), mouse monoclonal anti-GST (Abcam, ab92, dilution for WB 1:20000), mouse monoclonal anti-MBP (New England Biolabs, E8032, dilution for WB 1:10000). Peroxidase and Alexa-fluor conjugated secondary antibodies were purchased from Jackson Laboratories and Thermo, respectively. GFP-Booster Alexa Fluor® 488 (ChromoTek) was used in IF experiments.

### GFP-Trap pull down assay

For GFP pull down assay, HeLa cells expressing GFP::CIT-K constructs were transfected with an siRNA targeting the 3’UTR of *CIT-K* and synchronized (120 min after nocodazole release) as described above. Cells were then collected, washed in PBS, frozen in dry ice and stored at −80°C. Cell pellets were resuspended in 0.5 ml of lysis buffer (20 mM Tris-HCl, 150 mM NaCl, 2 mM MgCl_2_, 1 mM EGTA, 0.1% [v/v] NP-40, 1 mM DTT, 5% [v/v] glycerol, Roche Complete Protease Inhibitors and PhosSTOP Protein Phosphatase Inhibitors) and homogenized using a high-performance disperser (Fisher). The homogenate was clarified by centrifugation at 750 *g* for 15 min at 4°C and the supernatant was incubated with 40 μl of GFP-Trap magnetic Beads (ChromoTek) for 4 h on a rotating wheel at 4°C. Beads were then washed four times in 1 ml of lysis buffer for 5 min on a rotating wheel at 4°C, transferred to a new tube and washed one more time in 1 ml of PBS. After removing as much liquid as possible, beads were resuspended in 2x Laemmli sample buffer (Sigma Aldrich), boiled for 5 min and stored at −20°C. Proteins were separated by SDS PAGE, transferred onto PVDF membrane, and probed with antibodies to detect antigens.

### *In vitro* GST pull down assays

DNA fragments coding for CIT-K regions were generated by PCR and cloned into either pDEST15 (Thermo Fisher) to express N-terminal GST-tagged polypeptides or pKM596 (Fox et al., 2003) to express N-terminal MBP-tagged polypeptides *in E. coli*. DNA fragments coding for MKLP1_620-858_ and KIF14_901-1648_ were generated by PCR and cloned into pDEST15 (Thermo Fisher) to express N-terminal GST tagged polypeptides in *E. coli*. GST-tagged products were then purified using Glutathione Sepharose 4B according to manufacturer’s instruction (GE Healthcare). MBP-tagged products were purified on amylose resin according to manufacturer’s instruction (New England Biolabs). The bacteria pellet was resuspended in lysis buffer (20 mM Tris pH 7.5, 250 mM NaCl, 10 % glycerol, 0.2 % NP-40, 5 mM EDTA pH, 5 mM DTT, and a cocktail of Roche Complete Protease Inhibitors), sonicated, incubated for 20 minutes at 4°C, and then centrifuged. Equal volume of lysis buffer with 4% triton X-100 was then added to the supernatant, which was then loaded on amylose resin beads (New England Bioscience). Beads were washed five times with washing buffer (50 mM Tris-HCl pH 7.5, 250 mM NaCl, 0.2% [v/v] NP-40, 5 mM DTT) and eluted with 20 mM maltose.

For GST pull downs, MBP::CC1 fragments purified and eluted from beads were mixed with 25 μl of Glutathione Sepharose beads containing purified GST-tagged KIF14_901-1648_ or GST::MKLP1_620-856_ proteins. Samples were incubated in 300 μl of NET-N+ buffer (50mM Tris-HCl, pH 7.4, 150mM NaCl, 5mM EDTA, 0.2% NP-40, and a cocktail of Roche Complete Protease Inhibitors) for 60 minutes at 4°C on a rotating wheel and then washed 5 times with 500 μl of wash buffer (50mM Tris-HCl, pH 7.4, 250mM NaCl, 5mM EDTA, 0.2% NP-40, and a cocktail of Roche Complete Protease Inhibitors) followed by centrifugation at 500 *g* for 1 minute. Beads were resuspended in 25 μl of Laemmli SDS-PAGE sample buffer and analyzed by Western blot.

### Computational and statistical analysis

We used AlphaFold2 (2.3.2) (Jumper et al., 2021) to predict the structure of the longest and only CIT-K isoform that contains S699 (UniProt ID O14578-4).

Prism 10 (GraphPad) and Excel (Microsoft) were used for statistical analyses and to prepare graphs.

## Supporting information

This PDF file includes: Supplementary Figure S1, S2, S3, S4, S5, S6, S7 and associate legends

## Acknowledgments

We would like to thank Dr Marcos Rubio-Alarcón (University of Cambridge) for helping with AlphaFold2 and Dr Marco Geymonat (University of Cambridge) for helping with the purification of MBP-tagged proteins. This research was funded by two BBSRC UKRI grants to PPD (refs: BB/R001227/1 and BB/W01372X/1). EFJH was supported by a BBSRC DTP studentship.

