## Supplementary material for "Coordinated regulation of Citron kinase by CDK1 and Aurora B regulates midbody formation and stability": This PDF file includes: Supplementary Figure S1, S2, S3, S4, S5, S6, S7 and associate legends

**This PDF file includes:**
Supplementary Figure S1, S2, S3, S4, S5, S6, S7 and associate legends

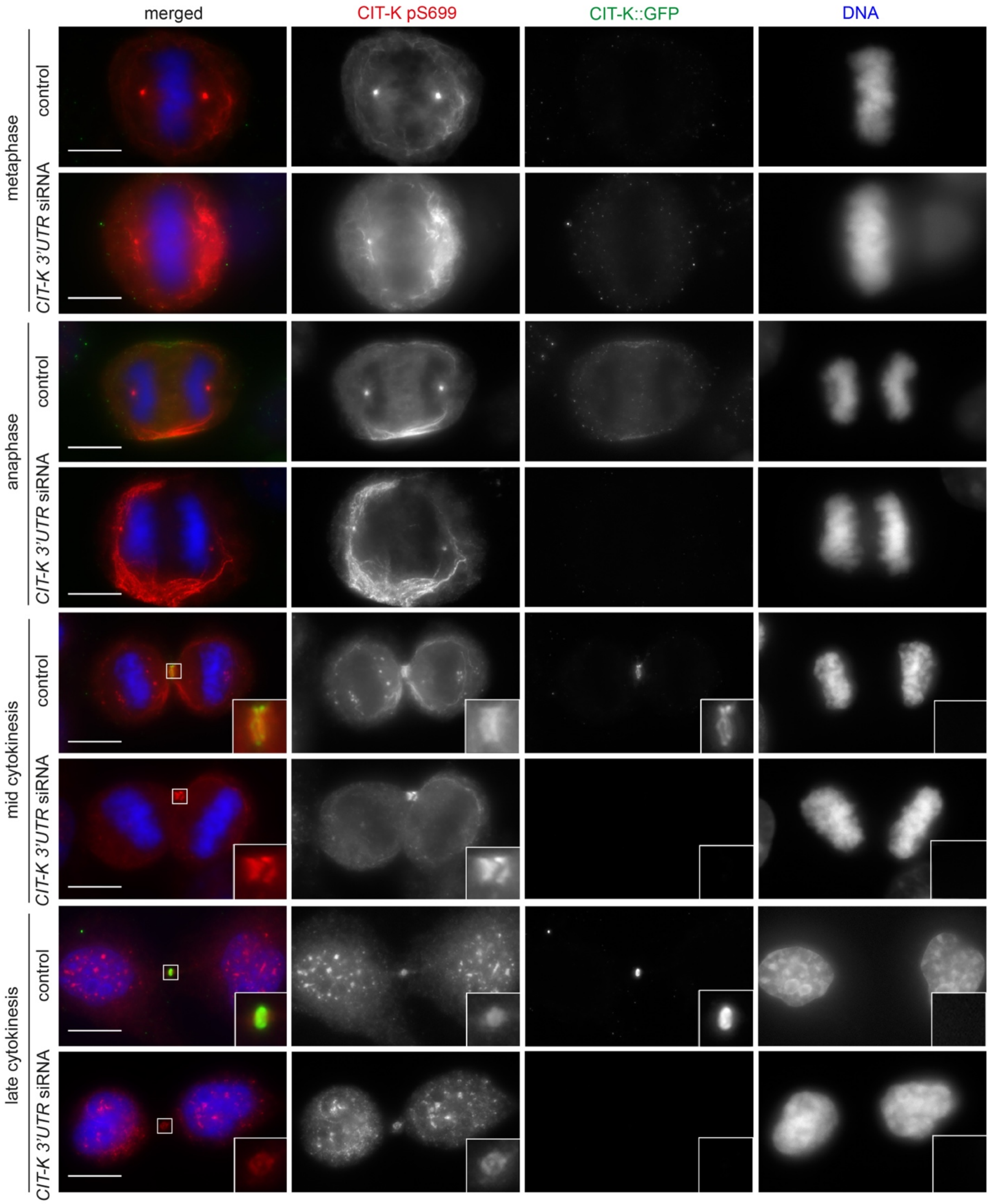


**Figure S1**. **The CIT-K pS699 antibody does not detect a specific signal in immunofluorescence experiments**. HeLa Kyoto cells stably expressing GFP-tagged CIT-K were treated with siRNAs directed against either a random sequence (control) or the 3'UTR of *CIT-K* and after 48 hours were fixed and stained to detect CIT-K::GFP (green), CIT-K pS699 (red) and tubulin (blue). The shape and thickness of microtubule bundles at the intercellular bridge were used as criteria to stage telophase cells. Insets show a 3X magnification of the midbody. Bars, 10 µm.

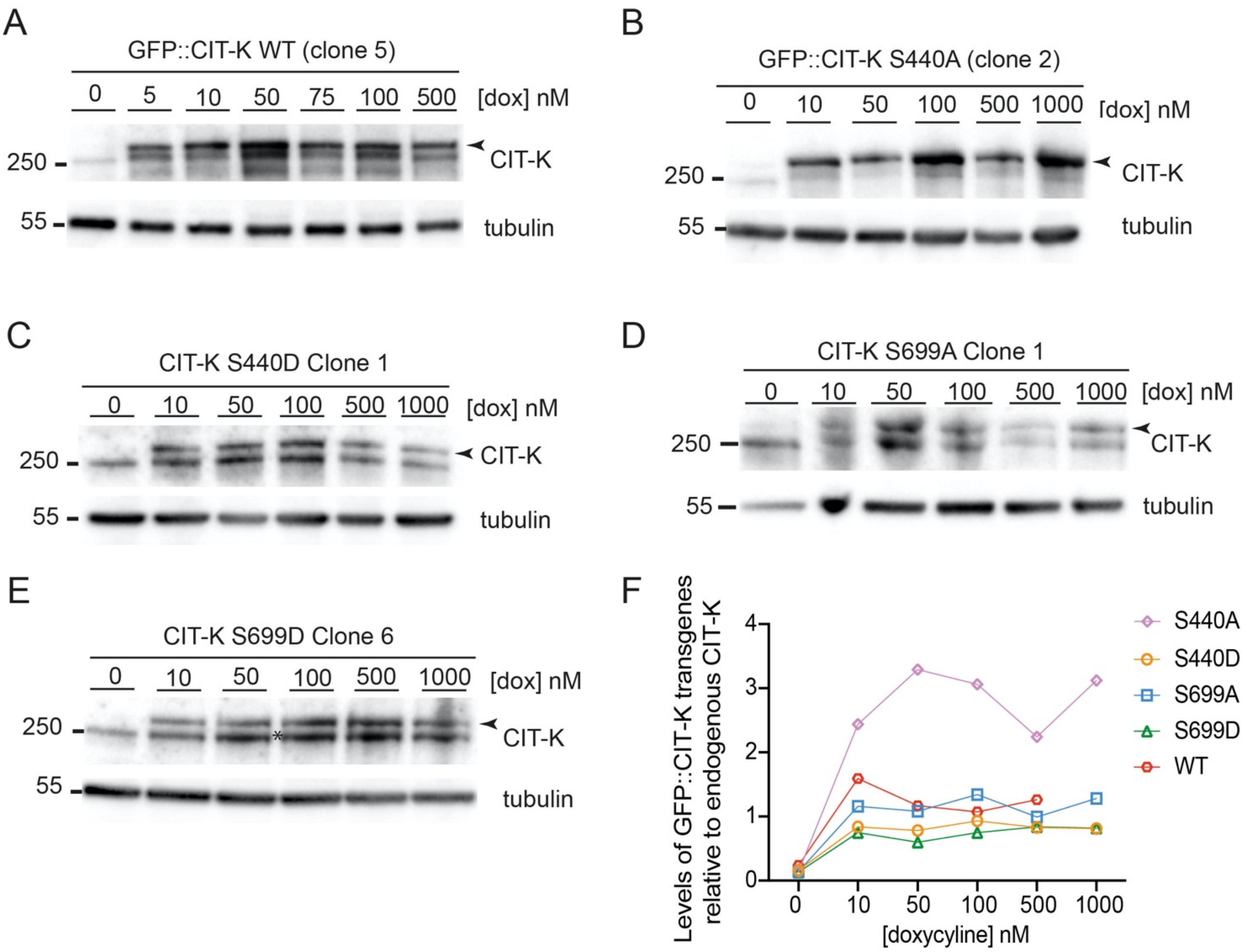


**Figure S2**. **Analysis of the level of expression of doxycycline-inducible GFP::CIT-K transgenes**. (A-E) Clones of cell lines stably expressing doxycycline-inducible phospho-dead and phosphomimetic S440 and S699 GFP::CIT-K transgenes were incubated with increasing concentrations (10-1000 nM) of doxycycline (dox). Proteins were extracted and analyzed by Western blot to detect GFP::CIT-K transgenes (indicated by arrowheads), endogenous untagged CIT-K, and tubulin as loading control. The numbers on the left indicate the sizes in kDa of the molecular mass marker. (F) Graph showing the quantification of the protein levels of GFP::CIT-K transgenes, normalized to tubulin and relative to the endogenous CIT-K.

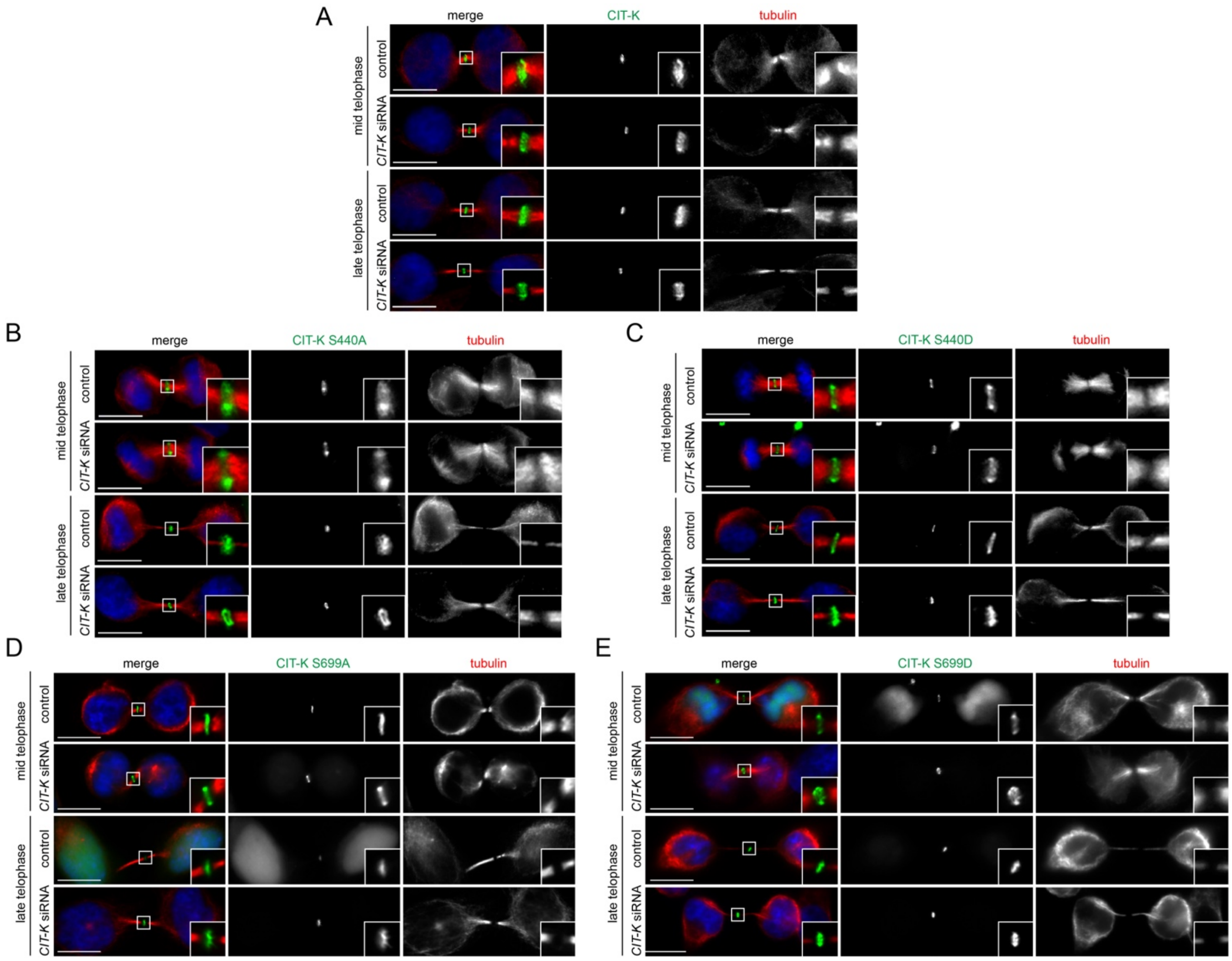


**Figure S3**. **Distribution of S440 and S699 phospho-mutant GFP::CIT-K proteins during cytokinesis**. (A-E) HeLa cells expressing wild-type (A) and phospho mutant (B-E) GFP::CIT-K constructs were treated with siRNAs directed against either a random sequence (control) or the 3'UTR of *CIT-K* and after 48 hours were fixed and stained to detect GFP::CIT-K (green), tubulin (red) and DNA (blue). The shape and thickness of microtubule bundles at the intercellular bridge were used as criteria to stage cells. Bars, 10 µm.

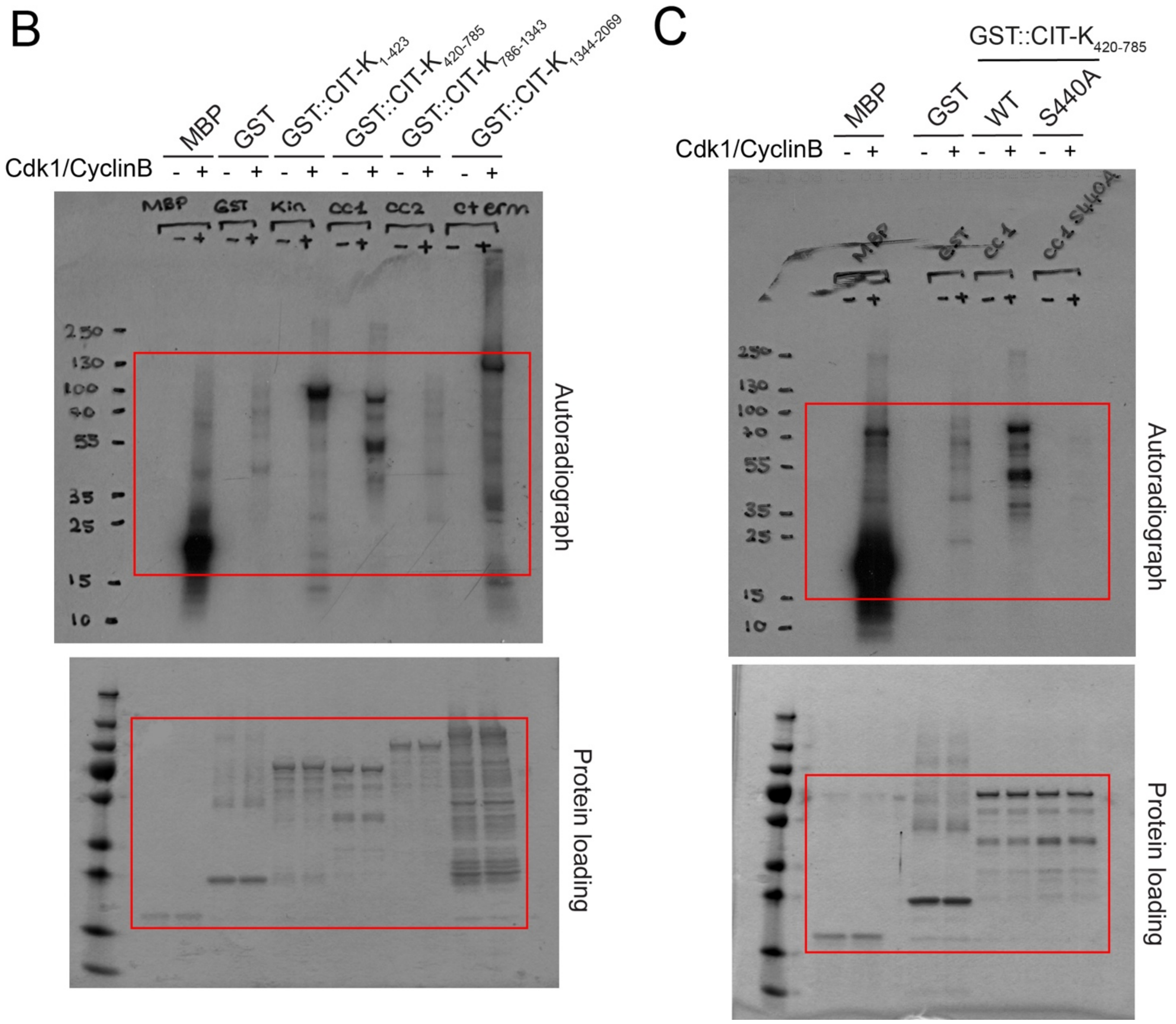


**Figure S4. Uncropped images of the autoradiographs and protein staining of the kinase assays shown in Figure 1.** The cropped sections shown in Figure 1 are marked by red rectangles.

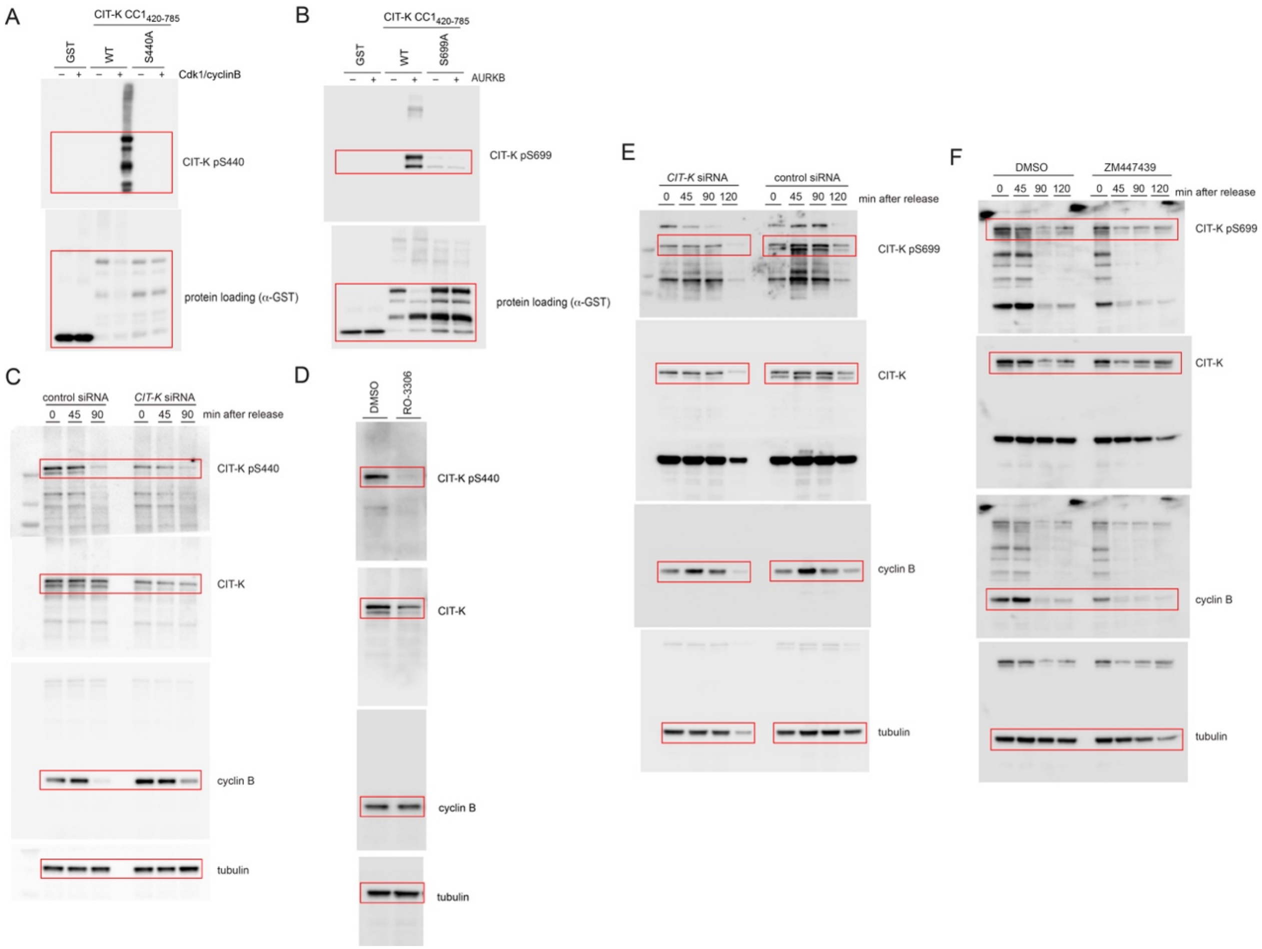


**Figure S5. Uncropped images of the Western blots shown in Figure 2.** The cropped sections shown in Figure 2 are marked by red rectangles.

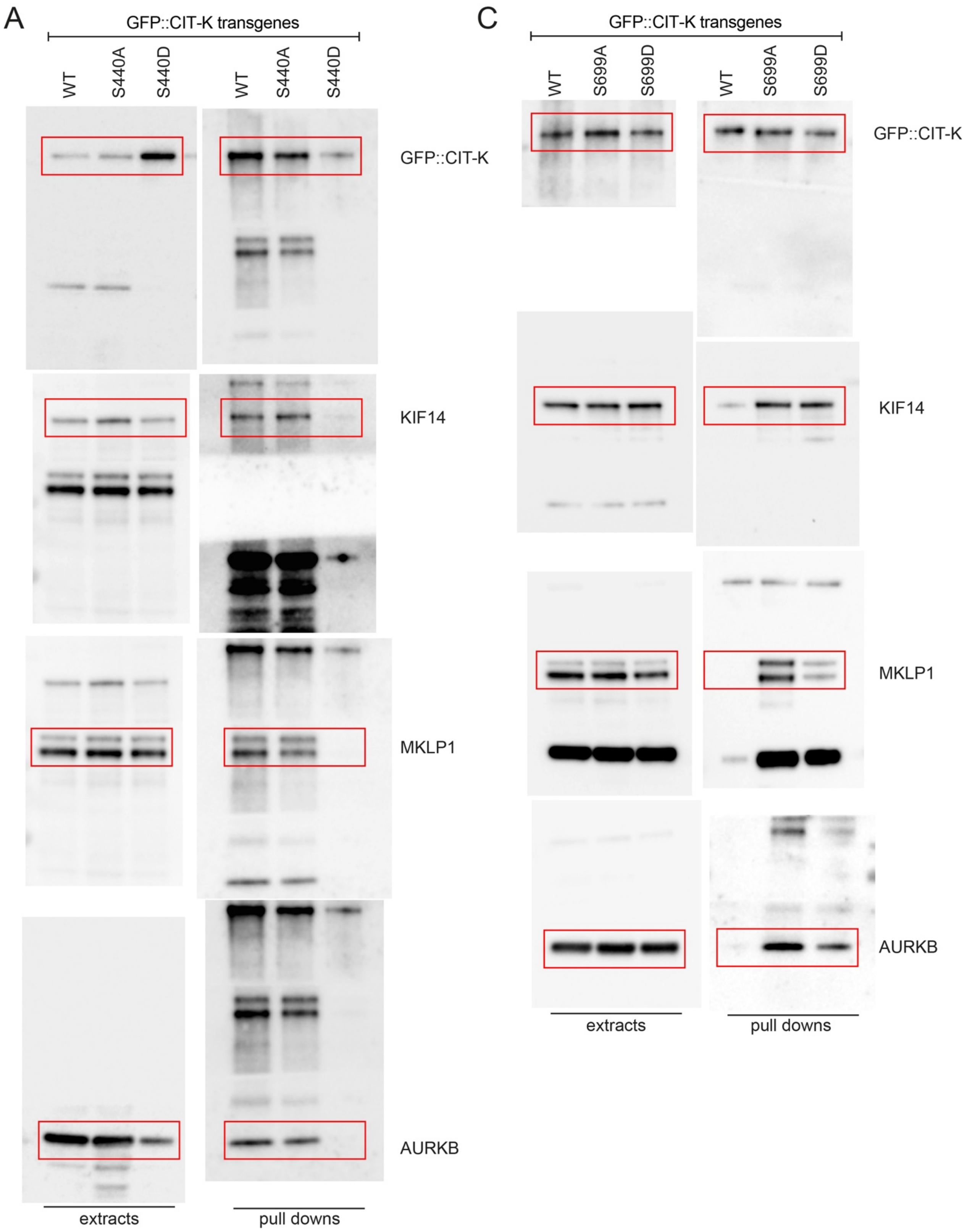


**Figure S6. Uncropped images of the Western blots shown in Figure 5.** The cropped sections shown in Figure 5 are marked by red rectangles.

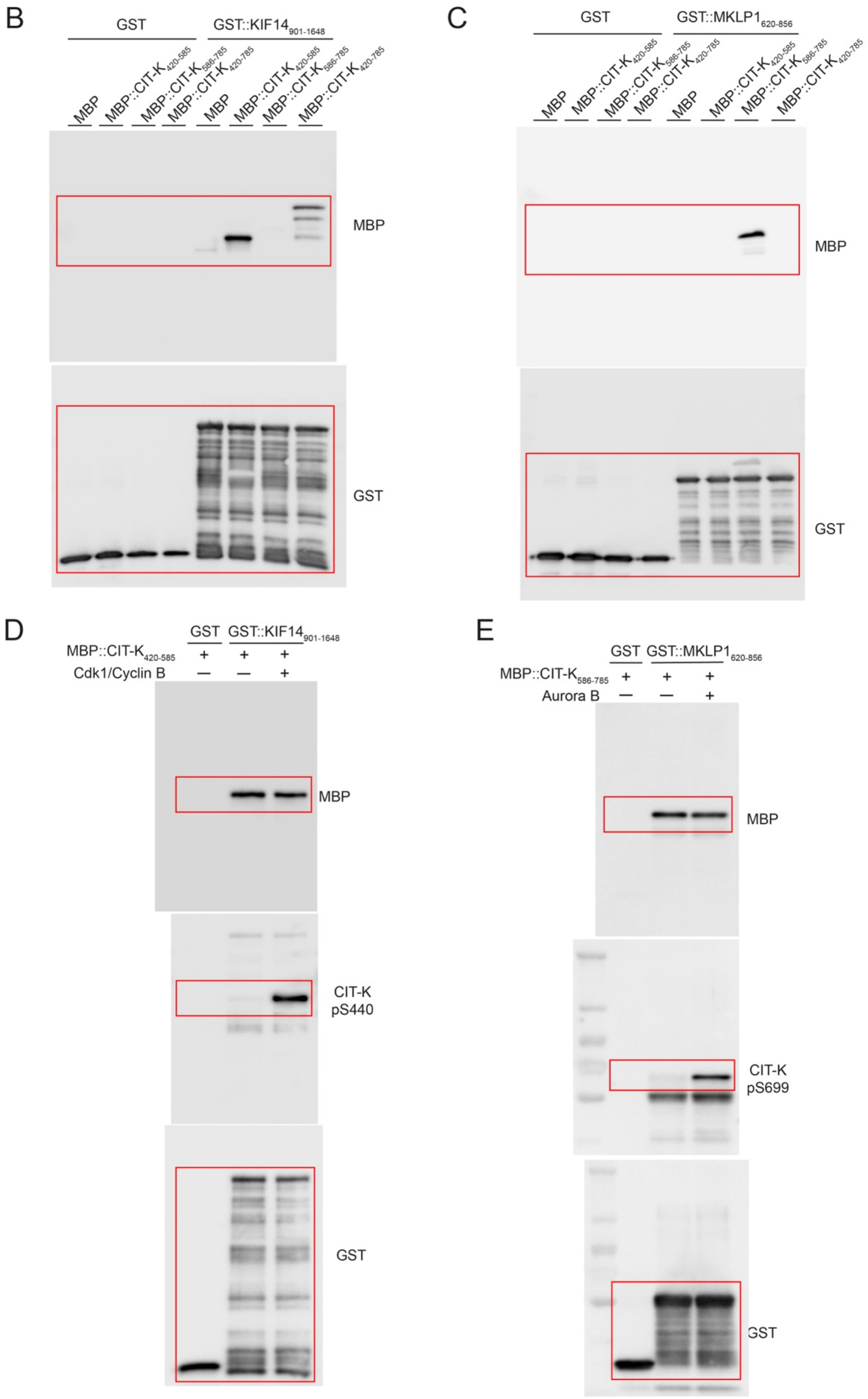


**Figure S7. Uncropped images of the Western blots shown in Figure 6.** The cropped sections shown in Figure 6 are marked by red rectangles.
